# Complete sequences of epidermin and nukacin encoding plasmids from oral-derived *Staphylococcus epidermidis* and their antibacterial activity

**DOI:** 10.1101/2021.09.24.461659

**Authors:** Kenta Nakazono, Mi Nguyen-Tra Le, Miki Kawada-Matsuo, Noy Kimheang, Junzo Hisatsune, Yuichi Oogai, Masanobu Nakata, Norifumi Nakamura, Motoyuki Sugai, Hitoshi Komatsuzawa

## Abstract

*Staphylococcus epidermidis* is a commensal bacterium in humans. To persist in the bacterial flora of the host, some bacteria produce antibacterial factors such as the antimicrobial peptides known as bacteriocins. In this study, we tried to isolate bacteriocin-producing *S. epidermidis* strains. Among 150 *S. epidermidis* isolates from the oral cavities of 287 volunteers, we detected two bacteriocin-producing strains, KSE56 and KSE650. Complete genome sequences of the two strains confirmed that they carried the epidermin-harbouring plasmid pEpi56 and the nukacin IVK45-like- harbouring plasmid pNuk650. The amino acid sequence of epidermin from KSE56 was identical to the previously reported sequence, but the epidermin synthesis-related genes were partially different. The prepeptide amino acid sequences of nukacin KSE650 and nukacin IVK45 showed one mismatch, but both mature peptides were entirely similar. pNuk650 was larger and had an additional seven ORFs compared to pIVK45. We then investigated the antibacterial activity of the two strains against several skin and oral bacteria and found their different activity patterns. In conclusion, we report the complete sequences of 2 plasmids coding for bacteriocins from *S. epidermidis*, which were partially different from those previously reported. Furthermore, this is the first report to show the complete sequence of an epidermin-carrying plasmid, pEpi56.

## Introduction

Staphylococci are classified into two groups, *Staphylococcus aureus* and coagulase - negative staphylococci (CoNS) due to their clinical importance. CoNS are abundant colonizers on the skin and are considered to contribute to the maintenance of skin integrity and homeostasis [1–3]. CoNS assist in immune activity to prevent pathogen colonization by immune cell priming, cutaneous inducing antimicrobial peptides from the epithelium, and direct production of antibacterial factors such as phenol-soluble modulins (PSMs) and bacteriocins [4–6]. Therefore, the colonization of CoNS provides several benefits to the host. However, CoNS are commonly isolated in clinical cultures and considered to be major nosocomial pathogens in humans [7, 8]. CoNS are often isolated from blood and indwelling medical implants such as intravascular catheters and urinary catheters, leading to opportunistic infectious diseases. In addition, most clinical isolates of *Staphylococcus epidermidis* carry the genes encoding for antibiotic resistance and biofilm formation, which significantly challenge current antibiotic therapy [9, 10].

Among staphylococci, *S. epidermids* is a major commensal bacterium in humans, mainly localized in the skin and nasal cavity [2, 3]. To persist among the bacterial flora of the host, it is well known that some bacteria produce antibacterial factors such as the antimicrobial peptides known as bacteriocins, and hydrogen peroxide [11–15].

Previously, it was reported that *S. epidermidis* produced bacteriocins such as epidermin [16–18], Pep5 [18–20], epilancin K7 [18, 21], epilancin 15X [22, 23], epicidin 280 [24] and nukacin IVK45 [25] to counter other bacterial species in the skin flora. However, the whole-genome sequences of these bacteriocin-producing strains have not been well characterized. Only, the nucleotide sequence of the plasmid coding for nukacin IVK45 was determined [25]. Bacteriocins are ribosomally synthesized and these *S. epidermidis* bacteriocins are classified as lantibiotics, which contain unusual amino acids such as lanthionine, β-methyllanthionine and dehydrated amino acids [11–13]. The antibacterial activity of these bacteriocins was characterized, but the main focus was on their effect against skin commensal bacteria. Since *S. epidermidis* is also found in the oral cavity [26, 27], it is also important to understand its antibacterial activity against oral bacteria. In this study, we isolated 135 *S. epidermidis* strains from the oral cavity and found 2 strains that produced epidermin and nukacin IVK45. We performed complete genome analysis of these 2 strains and identified the plasmids harbouring the epidermin or nukacin IVK45-like bacteriocin gene cluster. The nucleotide sequences of these plasmids were not entirely similar to the previously reported sequences. Additionally, we evaluated the antibacterial activity of these 2 bacteriocins against skin and oral commensal bacteria.

## Materials and methods

### Bacterial strains and growth conditions

*S. epidermidis* clinical isolates were grown in trypticase soy broth (TSB) (Becton, Dickinson and Company [BD], Franklin Lakes, NJ, USA) at 37°C. The *Staphylococcus aureus* MW2 strain and 14 sets of-inactivated mutants of each two-component system (TCS) were obtained previously [28]. Other bacteria used in this study are listed in Table 1. Staphylococcal strains and *Micrococcus luteus* were grown in TSB at 37°C and 30°C, respectively. Streptococcal strains were grown in TSB at 37°C with 5% CO2. *Cutibacterium acnes* was grown on sheep blood agar at 37℃ anaerobically. *Corynebacterium* and *Rothia mucilanginosa* were grown at 37°C in R medium and BHI (BD) aerobically, respectively. The composition of R medium is as follows: 1g of bacto peptone (BD), 0.5g of yeast extract (BD), 0.5g of malt extract (BD), 0.5g of casamino acids (BD), 0.2g of beef extract (BD), 0.2g of glycerol, 5mg of Tween 80, 0.1g of MgSO_4_ in 100 ml distilled water. When necessary, tetracycline (5 μg/ml) was added to the medium.

**Table 1.** Strains used in this study

| Strains | Character | Origin |
| --- | --- | --- |
| *Staphylococcus epidermidis* |  |  |
| KSE1 | Wild type | This study |
| KSE3 | Wild type | This study |
| KSE56 | Wild type | This study |
| KSE650 | Wild type | This study |
| KSE56- | KSE56 plasmid deleted | This study |
| KSE650- | KSE650 plasmid deleted | This study |
| *Staphylococcus warneri* ISK-1 | Wild type | [29] |
| *Staphylococcus hominis* JCM31912 | Wild type | Riken BRC<sup>1</sup> |
| *Staphylococcus haemolyticus* JCM2416 | Wild type | Riken BRC<sup>1</sup> |
| *Staphylococcus capitis* JCM2420 | Wild type | Riken BRC<sup>1</sup> |
| *Staphylococcus simulans* JCM2424 | Wild type | Riken BRC<sup>1</sup> |
| *Staphylococcus saprophyticus* JCM20595 | Wild type | Riken BRC<sup>1</sup> |
| *Cutibacterium acnes* JCM6425 | Wild type | Riken BRC<sup>1</sup> |
| *Corynebacterium accolens* JCM8331 | Wild type | Riken BRC<sup>1</sup> |
| *Corynebacterium pseudodiphtheriticum* JCM1320 | Wild type | Riken BRC<sup>1</sup> |
| *Rothia mucilaginosa* JCM10910 | Wild type | Riken BRC<sup>1</sup> |
| *Micrococcus luteus* JCM1464 | Wild type | Riken BRC<sup>1</sup> |
| *Streptococcus mutans* UA159 | Wild type | [30] |
| *Streptococcus sanguinis* GTC217 | Wild type | Gifu University |
| *Streptococcus salivarius* GTC215 | Wild type | Gifu University |
| *Streptococcus gordonii* JCM12995 |  | Riken BRC<sup>1</sup> |
| *Staphylococcus aureus* COL | Wild type | [31] |
| RN4220 | NCTS8325 derivative | [32] |
| MW2 | clinical strain, methicillin-resistant (*mecA*+) ) | [33] |
| Δ TCS2 | *MW0918-99* inactivation in MW2, Tc<sup>r</sup> <sup>2</sup> | [28] |
| Δ TCS3 | *lytSR* inactivation in MW2, Tc<sup>r</sup> | [28] |
| Δ TCS4 | *apsRS* inactivation in MW2, Tc<sup>r</sup> | [28] |
| Δ TCS5 | *saeRS* inactivation in MW2, Tc<sup>r</sup> | [28] |
| Δ TCS6 | *MW1208-09* inactivation in MW2, Tc<sup>r</sup> | [28] |
| Δ TCS7 | *arlRS* inactivation in MW2, Tc<sup>r</sup> | [28] |
| Δ TCS8 | *srrAB* inactivation in MW2, Tc<sup>r</sup> | [28] |
| Δ TCS9 | *phoPR* inactivation in MW2, Tc<sup>r</sup> | [28] |
| Δ TCS10 | *MW1789-90* inactivation in MW2, Tc<sup>r</sup> | [28] |
| Δ TCS11 | *vraSR* inactivation in MW2, Tc<sup>r</sup> | [28] |
| Δ TCS12 | *agrCA* inactivation in MW2, Tc<sup>r</sup> | [28] |
| Δ TCS13 | *kdpDE* inactivation in MW2, Tc<sup>r</sup> | [28] |
| Δ TCS14 | *hssRS* inactivation in MW2, Tc<sup>r</sup> | [28] |
| Δ TCS15 | *nreBC* inactivation in MW2, Tc<sup>r</sup> | [28] |
| Δ TCS16 | *braRS* inactivation in MW2, Tc<sup>r</sup> | [28] |

1. Japan Collection of Microorganisms
2. Tetracycline resistance

### Isolation of *Staphylococcus epidermidis* from the oral cavity

*S. epidermidis* strains were isolated from the oral cavities of 287 volunteers. Saliva collected from the oral cavity was plated on No.110 medium (Eiken Chemical Co. Ltd, Tokyo, Japan) and incubated for 2 days at 37°C. The strains were picked from a single white colony on the agar and further investigated by PCR with specific primers for *S. epidermidis* (forward primer: GGCAAATTTGTGGGTCAAGA, reverse primer: TGGCTAATGGTTTGTCACCA). Isolated *S. epidermidis* strains were replated on TSB containing 2% agar (TSA) medium. The isolated strains were then replated again on TSA to pick up a single colony and finally, *S. epidermidis* confirmed by PCR was used in this study. Clinical isolates were designated as KSE strains. *S. epidermidis* isolation was approved by the ethics committee of the Kagoshima University Graduate School of Medical and Dental Sciences (No. 701) and the Ethical Committee for Epidemiology of Hiroshima University (E-1998). All methods were performed in accordance with the approved guidelines and regulations.

### Screening of bacteriocin producing *S. epidermidis*

To investigate bacteriocin production among *S. epidermidis* strains, we performed a direct assay using *S. aureus* MW2 *braRS* knockout mutant as an indicator strain because this mutant showed increased susceptibility to several bacteriocins [34]. Overnight cultures of *S. epidermidis* strains were spotted on a TSA plate and cultured at 37°C for 24 h. Then, 3.5 ml of prewarmed half-strength TSB soft agar (1%) containing *braRS* knockout mutant cells (10^7^ cells/ml) were poured over the TSA plate. The plates were incubated at 37°C for 24 h. The strains which showed the growth inhibition zones surrounding *S. epidermidis* strain were picked up. The strains were reconfirmed for bacteriocin production by the direct assay again.

### Complete genome sequences of bacteriocin-producing *S. epidermidis* strains

To perform whole-genome sequencing of *S. epidermidis* strains, DNA was extracted from each strain. *S. epidermidis* cells grown overnight in 5 ml TSB were collected and then suspended in 0.5 ml of CS buffer (100 mM Tris-HCl [pH 7.5], 150 mM NaCl, 10 mM EDTA) containing lysostaphin (Sigma-Aldrich, St. Louis, MO, USA) (final concentration: 50 μg/ml) and RNase (Nippon Gene, Tokyo, Japan) (final: 20 μg/ml). After incubation at 37°C for 1 h, proteinase K (Nacalai Tesque, Kyoto, Japan) (final: 150 μg/ml) and SDS (final 1%) were added, followed by incubation at 55°C for 5 h.

After treatment with phenol followed by phenol-chloroform, DNA was precipitated by ethanol. Whole-genome sequencing (WGS) of *S. epidermidis* strains **w**as performed using the Illumina MiSeq sequencing platform, followed by annotation with Rapid Annotation using Subsystem Technology (RAST) version 2.0 [35]. After confirming the presence of bacteriocin genes using WGS, long-read sequencing by MinION (Oxford Nanopore Technologies, UK) was carried out to determine the complete sequences of the chromosomes and plasmids of these strains. Hybrid assembly of Illumina short reads and MinION long reads was performed with Unicycler v0.4.8. The complete sequences of plasmids harbouring bacteriocin genes were selected, including epidermin-carrying plasmid pEpi56 and nukacin-carrying plasmid pNuk650. Each plasmid was compared with publicly available plasmids or gene clusters, including the *epiY’-epiP* gene cluster (X62386), *epiG-epiT’’* gene cluster (U77778), and pIVK45 (accession number KP702950).

## Accession numbers

The complete plasmids carrying epidermin (pEpi56) and nukacin (pNuk650) have been deposited in the NCBI database under accession numbers OK031036 and OK031035, respectively.

### Identification of epidermin and nukacin KSE650 produced by *S. epidermidis*

To identify the bacteriocin, we purified the bacteriocin from two *S. epidermidis* strains. Overnight cultures (500 ml) of *S. epidermidis* KSE56 and KSE650 were centrifuged at 4,000 x g for 15 min. Macro-Prep cationic resin (1.5 ml)(Bio rad, USA) was added to the supernatant and stirred for 12 h. The resin was collected into an open column, then washed three times with 10 ml of 25 mM ammonium acetate (pH 7.5). To elute the bacteriocin, the resin was treated with 500 μl of 5% acetic acid. This elution was repeated 10 times. After each fraction was evaporated completely, the samples were dissolved in 50 μl of distilled water. Each solution was tested for antibacterial activity against *M. luteus*. Overnight cultures of *M. luteus* (100 μl) were inoculated on TSA plates. Then, 5 μl of each solution was spotted on TSA. After overnight incubation at 37°C, growth inhibition was observed. Samples with antibacterial activity were subjected to HPLC chromatography using an Octadecyl C18 column. After equilibrating the column with 0.1% TFA water, the sample was injected. Thereafter, a linear gradient of 0 to 60% acetonitrile for 30 min was applied to the column. Each peak was fractionated, and the samples were evaporated, then dissolved with 50 μl of distilled water. Subsequently, the antibacterial activity of each fraction was tested with the method above. ESI-MS analysis was performed by LTQ Orbitrap XL (Thermo Fisher Scientific, USA).

### Isolation of the strain curing bacteriocin-encoded plasmid

Plasmid deletion in KSE56 and KSE650 was performed with the method described elsewhere [36]. Overnight cultures of KSE56 or KSE650 were inoculated into 5 ml of fresh TSB and incubated at 37°C with shaking. When the OD660 reached 0.5, acriflavine was added at a concentration of 25 μg/ml. After incubation for 12 h, the culture was diluted and plated on TSA. After 24 h of incubation at 37°C, colonies were picked, replated on TSA and then incubated at 37°C for 24 h. Next, 0.75% soft agar (3.5 ml) containing *Bacillus coagulans* (200 μl of overnight culture) was poured on that plate and incubated at 37°C for 24 h. The strains with no inhibitory zone were picked. Finally, PCR was performed using specific primers for *S. epidermidis-*specific genes and bacteriocin genes coding for nukacin KSE650 or epidermin.

### Direct assay

To evaluate the antibacterial activity of epidermin, nukacin KSE650 and nukacin ISK-1, a direct assay was performed with a previously described method [34]. An overnight culture of the bacteriocin-producing strain was spotted on a TSA plate and cultured at 37°C for 24 h. Then, 3.5 ml of prewarmed half-strength TSB soft agar (1%) containing indicator bacterial cells (10^7^ cells/ml) was poured over the TSA plate. The plates were incubated at 37°C for 16 h. The diameters of the growth inhibition zones surrounding the bacteriocin-producing strains were measured in three directions. Three independent experiments were performed, and the average diameter was calculated.

### Co-culture of S. epidermidis with M. luteus

For analysis of the proportion of each bacterium (*S. epidermidis* and *M. luteus*) in coculture by qPCR, we first set up the method for the calculation of bacterial cell number by qPCR. A single overnight culture of the bacterium was first adjusted to OD660=1.0, and then a 10-fold serial dilution was performed in 500 µl of lysis buffer. After heating at 95°C for 15 min, samples were centrifuged at 15,000 x rpm for 10 min. Using the supernatant, qPCR was performed with the respective specific primers. For *S. epidermidis*, the forward and reverse primers used were GGCAAATTTGTGGGTCAAGA and TGGCTAATGGTTTGTCACCA, respectively. For *M. luteus*, the forward and reverse primers were GGGTTGCGATACTGTGAGGT and TTCGGGTGTTACCGACTTTC, respectively.

Finally, the linear relationship between bacterial cell number and cut off value (Ct value) was constructed in each bacterium. Overnight cultures of *S. epidermidis* KSE1 (no bacteriocin production), KSE56, KSE650 and *M. luteus* were adjusted to OD660=1.0, and the bacterial culture was diluted to 10-fold. Next, 100 µl of *S. epidermidis* culture and *M. luteus* were mixed thoroughly. A small portion (20 μl) of mixed culture was spotted on TSA. After overnight incubation at 37°C, the bacterial colonies growing on agar plates were scraped and suspended in 500 µl of lysis buffer. After heating at 95°C for 15 min, the bacterial suspension was centrifuged at 15,000 x rpm for 10 min and the culture supernatant was stocked as the template for quantitative PCR (qPCR). qPCR was performed using appropriate specific primers to determine the cell number of each bacterium in the coculture samples. Finally, the proportion of 2 bacterial species was determined. Three independent experiments were performed. Post hoc multiple comparisons were made using Tukey’s test.

## Results

### Isolation of *S. epidermidis* that produced bacteriocin

From 287 volunteers, 150 *S. epidermidis* strains (52.3%) were isolated from the oral cavity. Among 150 *S. epidermidis* strains, 2 strains showing a clear inhibitory zone against the *S. aureus* MW2 *braRS* inactivated mutant were identified by the direct method (Fig.1).

**Figure 1.**
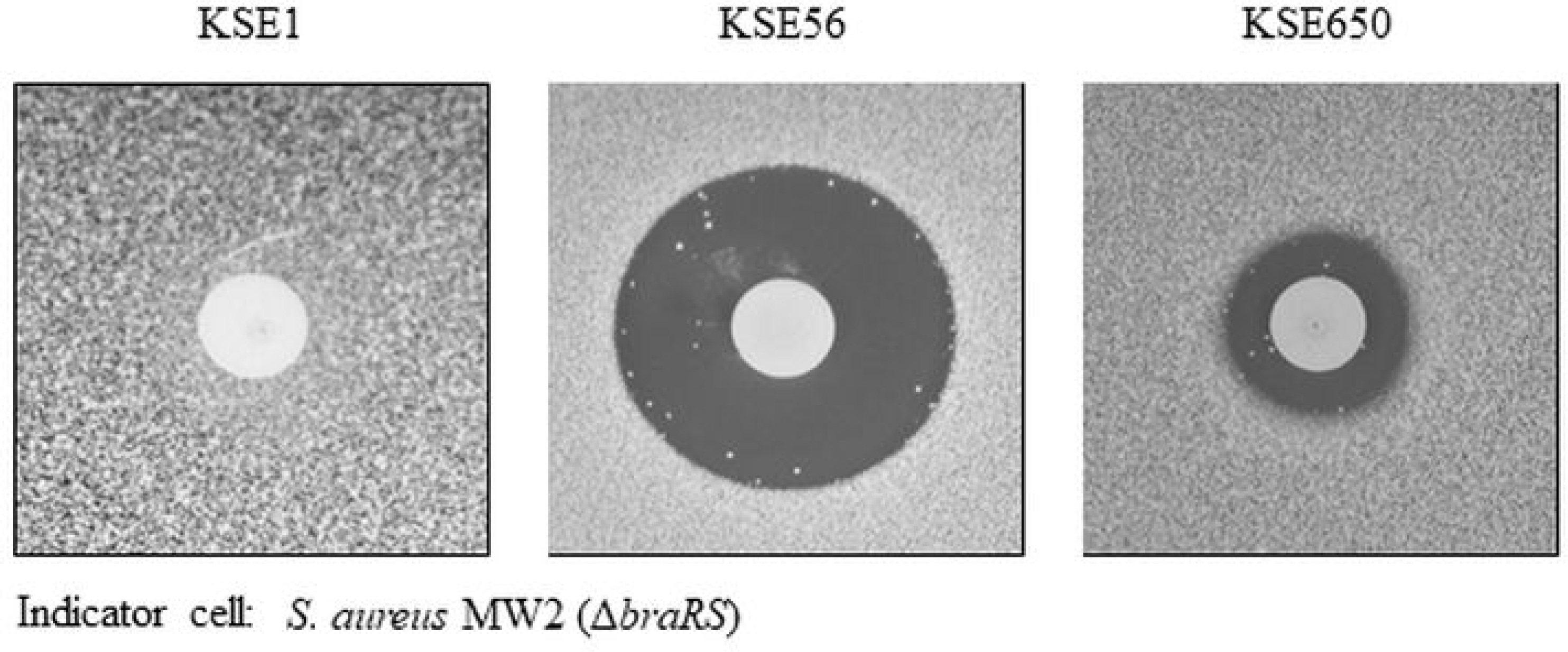
Direct assay of bacteriocin-producing *S. epidermidis* against *braRS*- inactivated *S. aureus*. The antibacterial activity of bacteriocin-producing *S. epidermidis* was evaluated by the direct assay using *S. aureus* MW2 *braRS*-inactivated mutant.

### Nucleotide sequence of epidermin-encoding plasmid

The size of the entire plasmid, pEpi56, is 64,386 bp, with 81 ORFs (Fig. 2a and Table 2). The plasmid contains epidermin synthesis genes (*epiA* coding for epidermin KSE56, modification genes *epiBCD*, processing genes *epiP*, export genes *epiHT*, immunity genes *epiGEF*, and regulatory gene *epiQ*), replication-related genes, and other genes including the genes coding for hypothetical proteins (Table 2). Compared with epidermin-related genes in the Tü3298 strain (16), *epiT*, which codes for an exporter, was intactin pEpi56, while a gene disrupted into two fragments (*epiT’* and *epiT’’* or *epiY and epiY’*) was found in the Tü3298 strain (Fig. 2b, Supplemental Fig. 1). The nucleotide sequence of *epiA* in KSE56 showed 2 mismatches with that of the Tü3298 strain (Supplemental Fig. 2). However, the amino acid sequence of epidermin KSE56 showed 100% identity with that in the Tü3298 strain.

**Figure 2.**
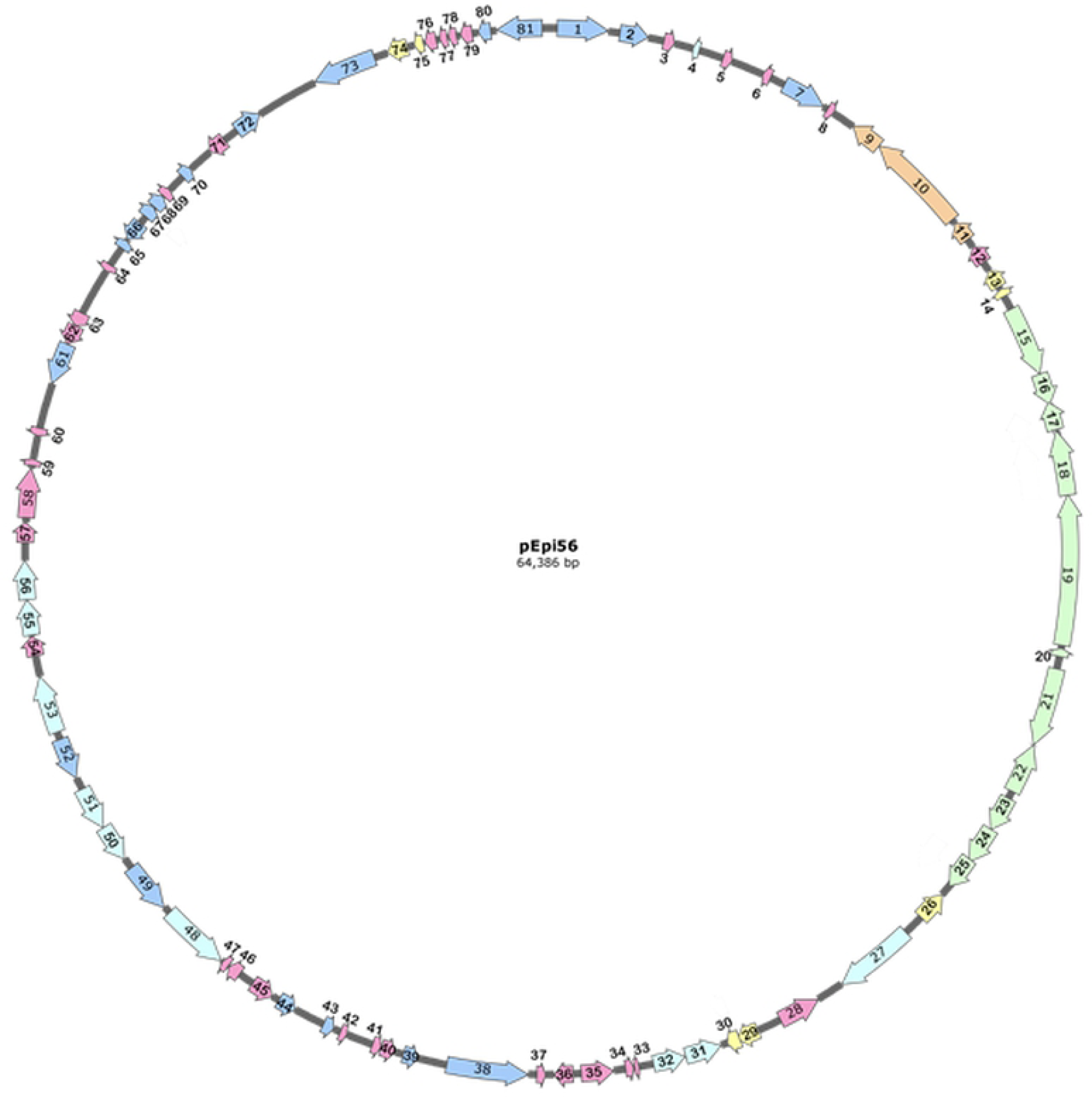
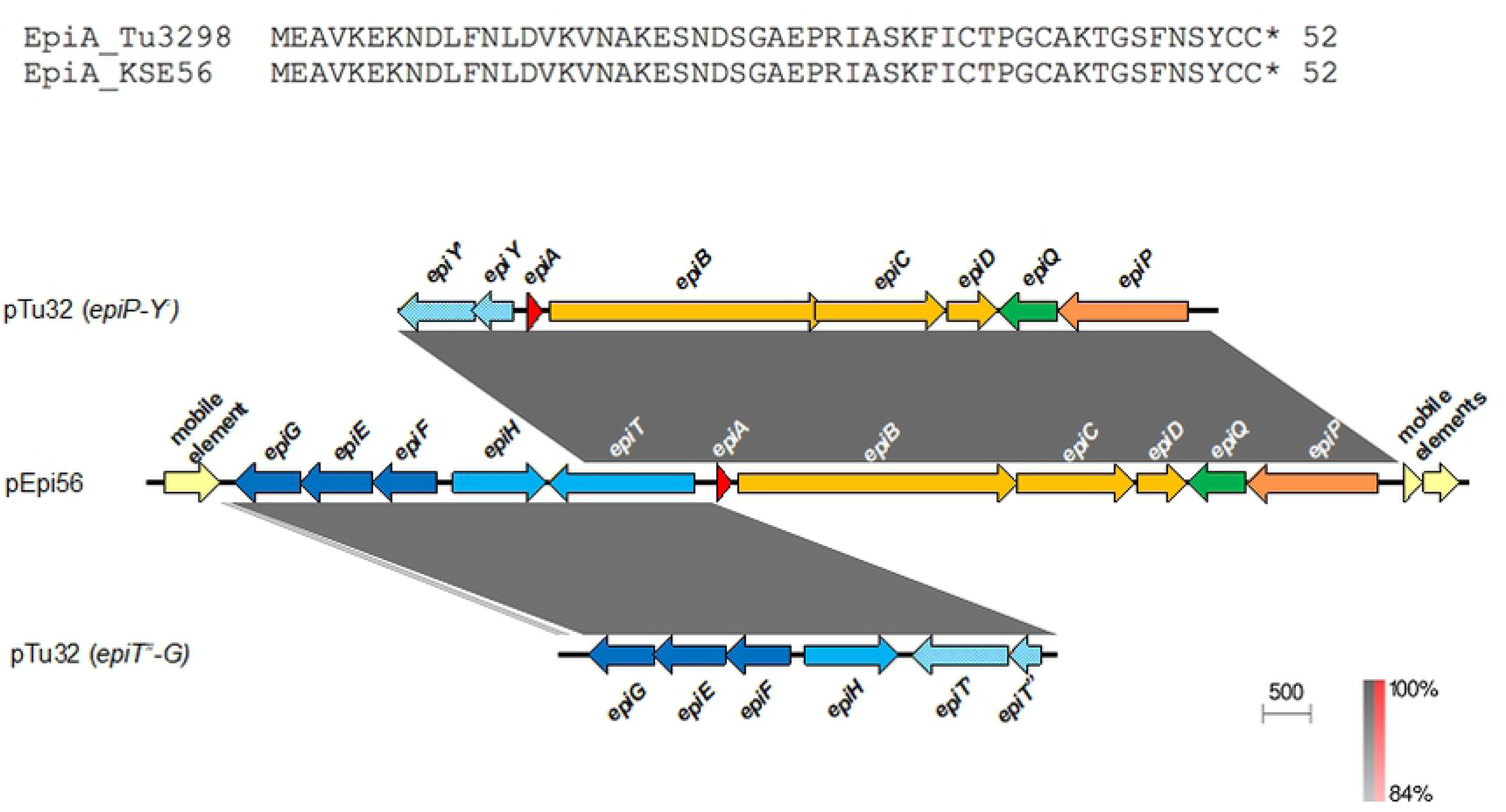
Gene map of the plasmid carrying epidermin in KSE56. (a) Epidermin-encoding plasmid from KSE56 (pEpi56). ORFs are shown as arrows, indicating the orientation of transcription. The arrow numbers indicate the ORF number displayed in Table 2. Colors indicate the classification of gene function. (b) Bacteriocin-coding region (KSE56 epidermin). The bacteriocin-coding region from pEpi56 was compared with pTu32 *epiP-Y’* (accession number X62386) and pTu32 *epiT”-G* (accession number U77778). Striped blue arrows indicate truncated *epiT*.

**Table 2.**
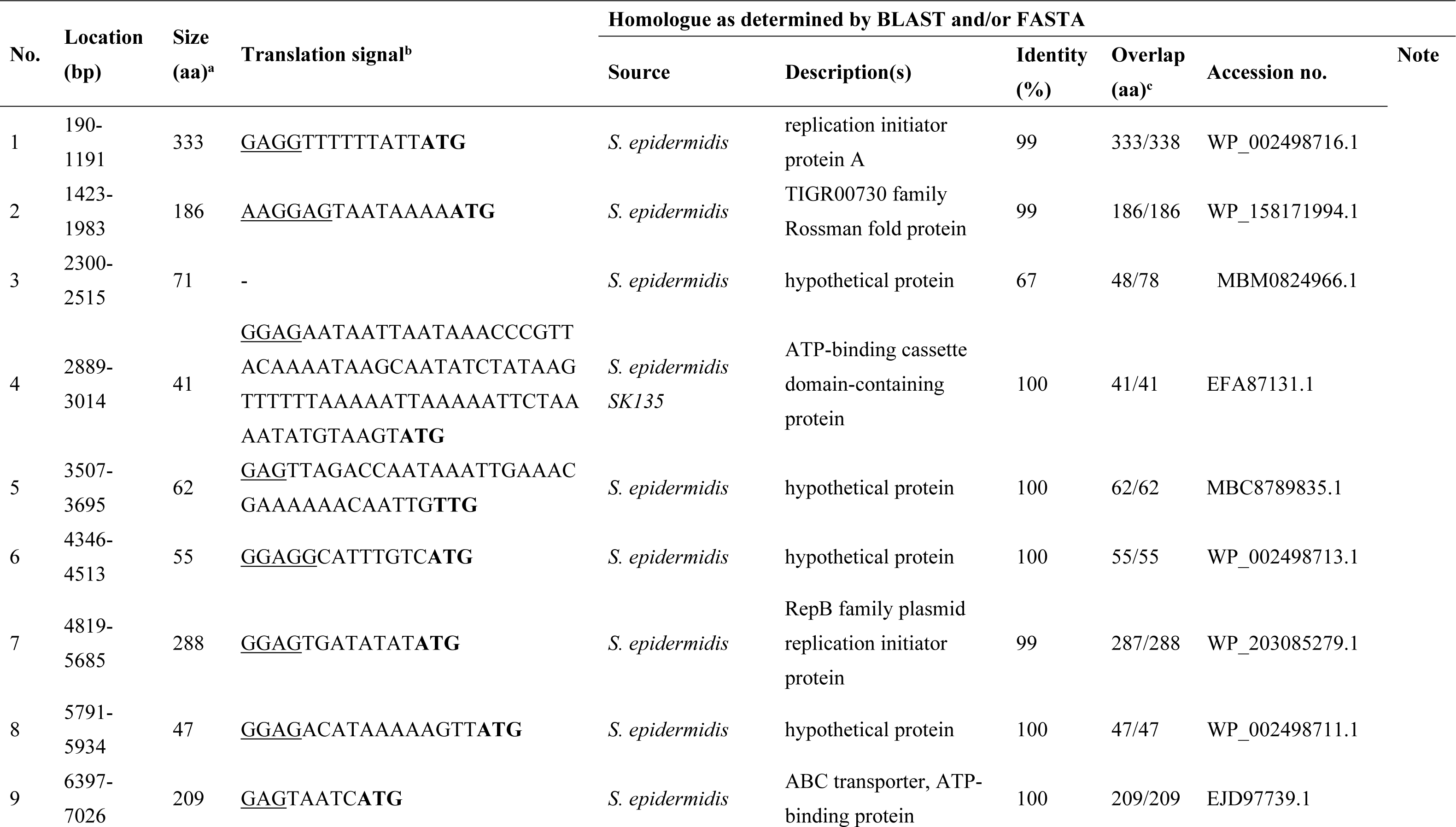
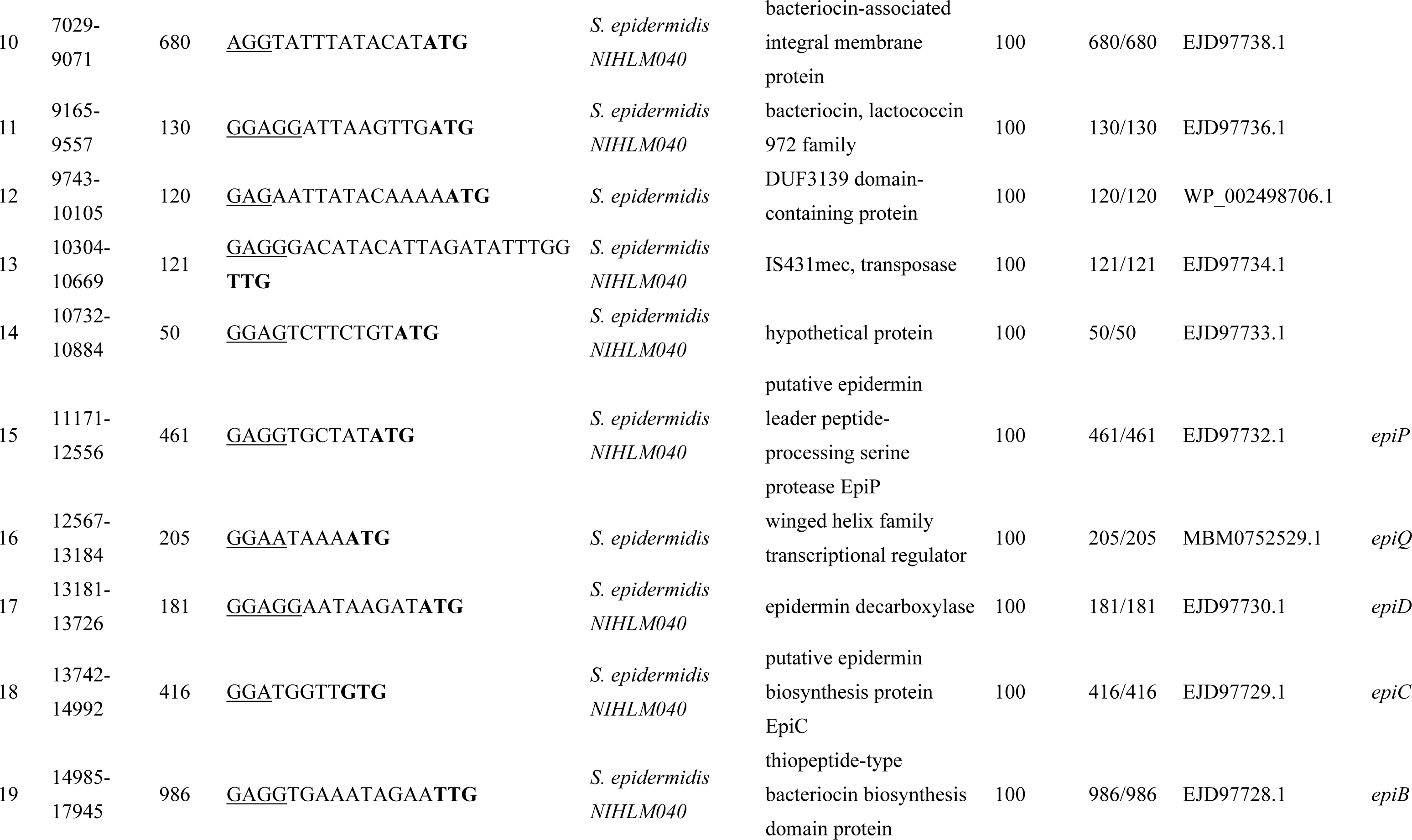
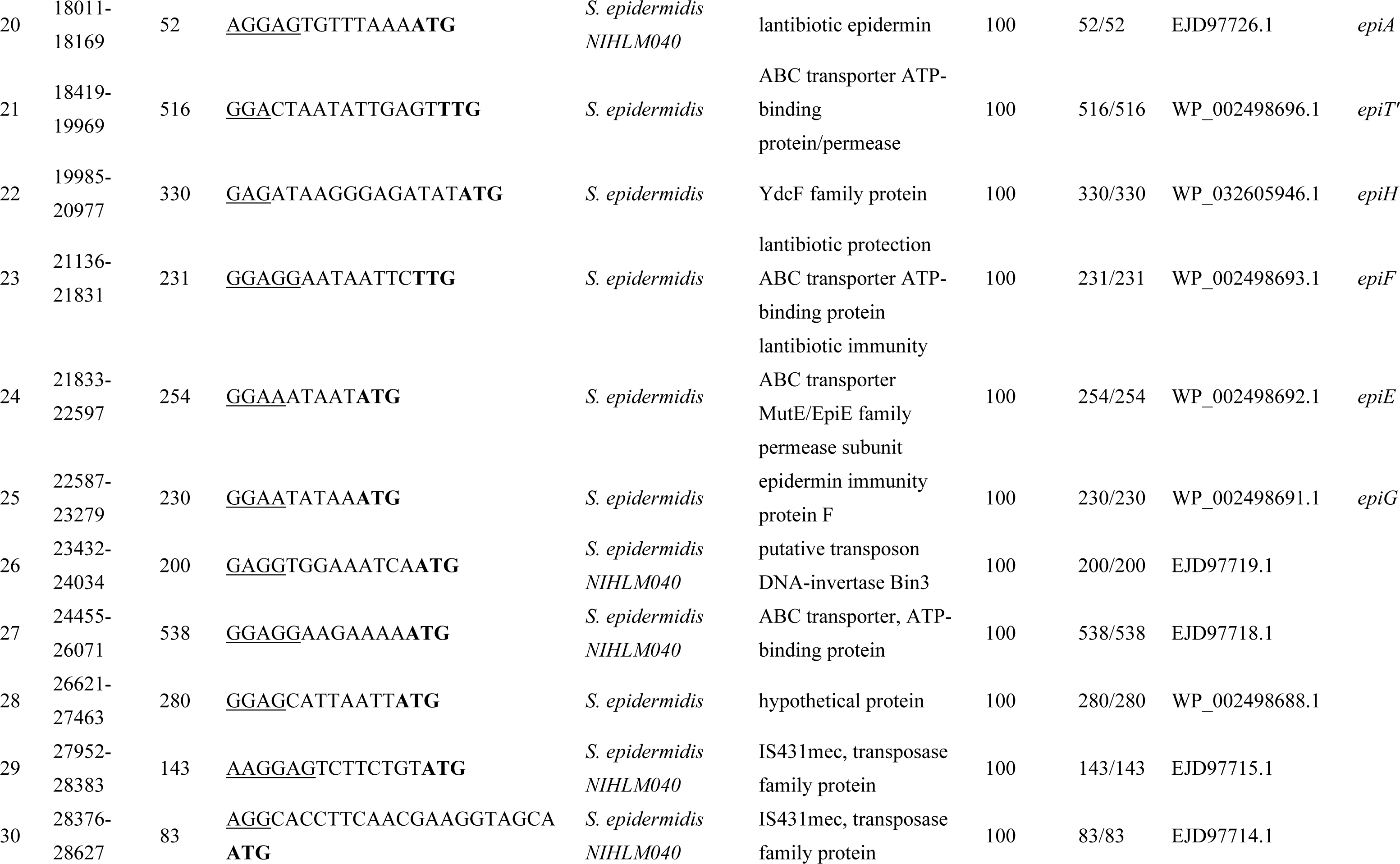
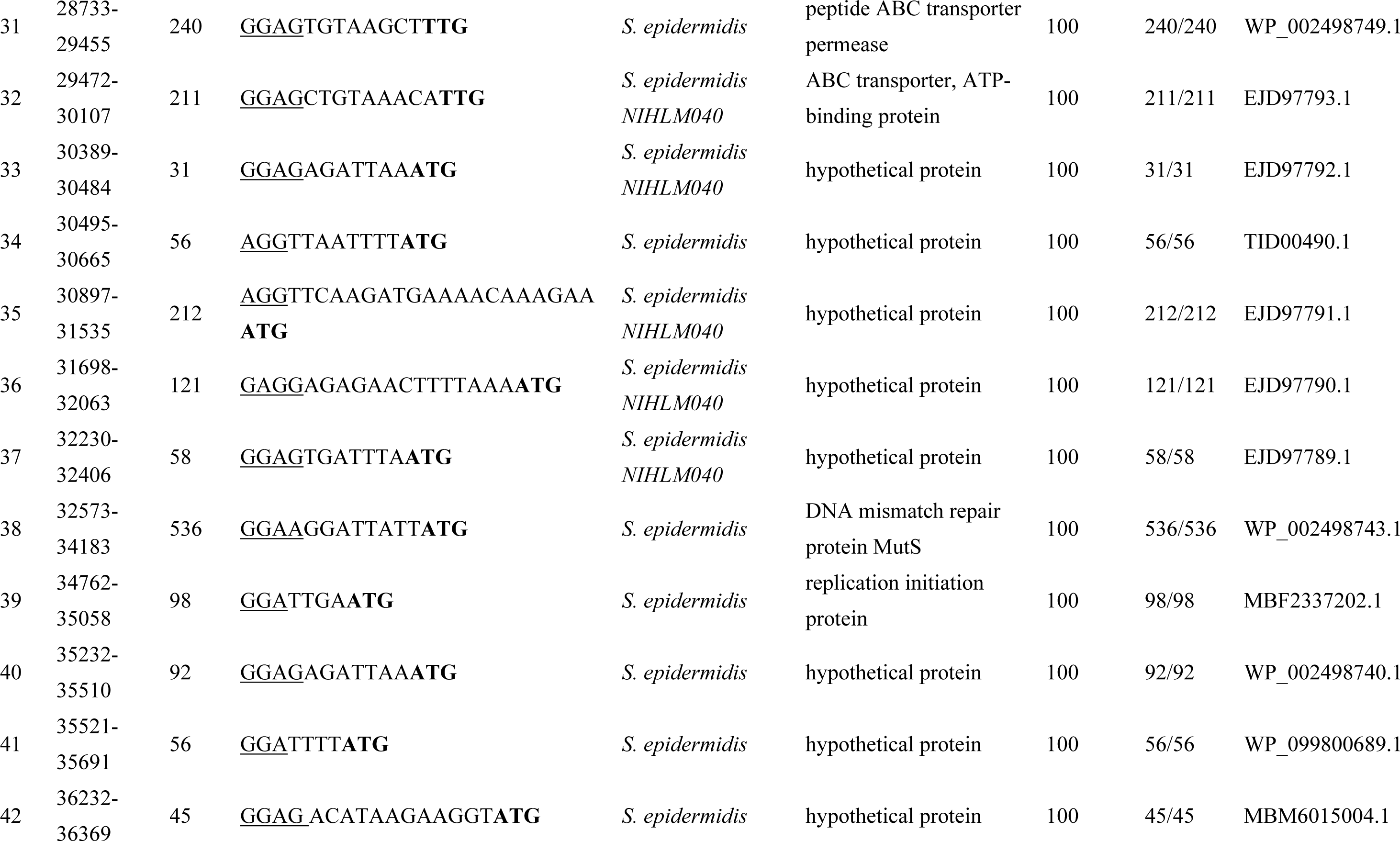
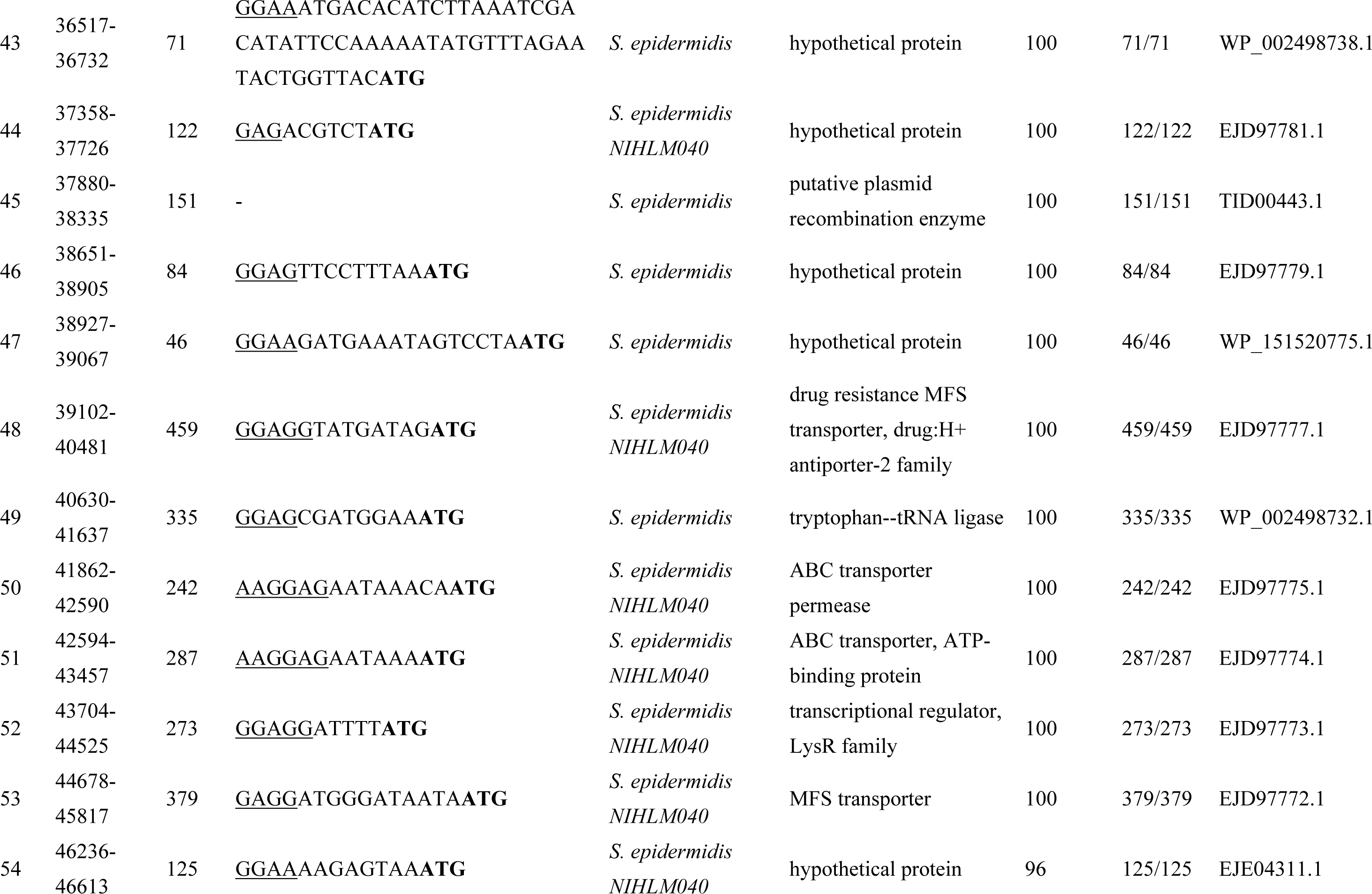
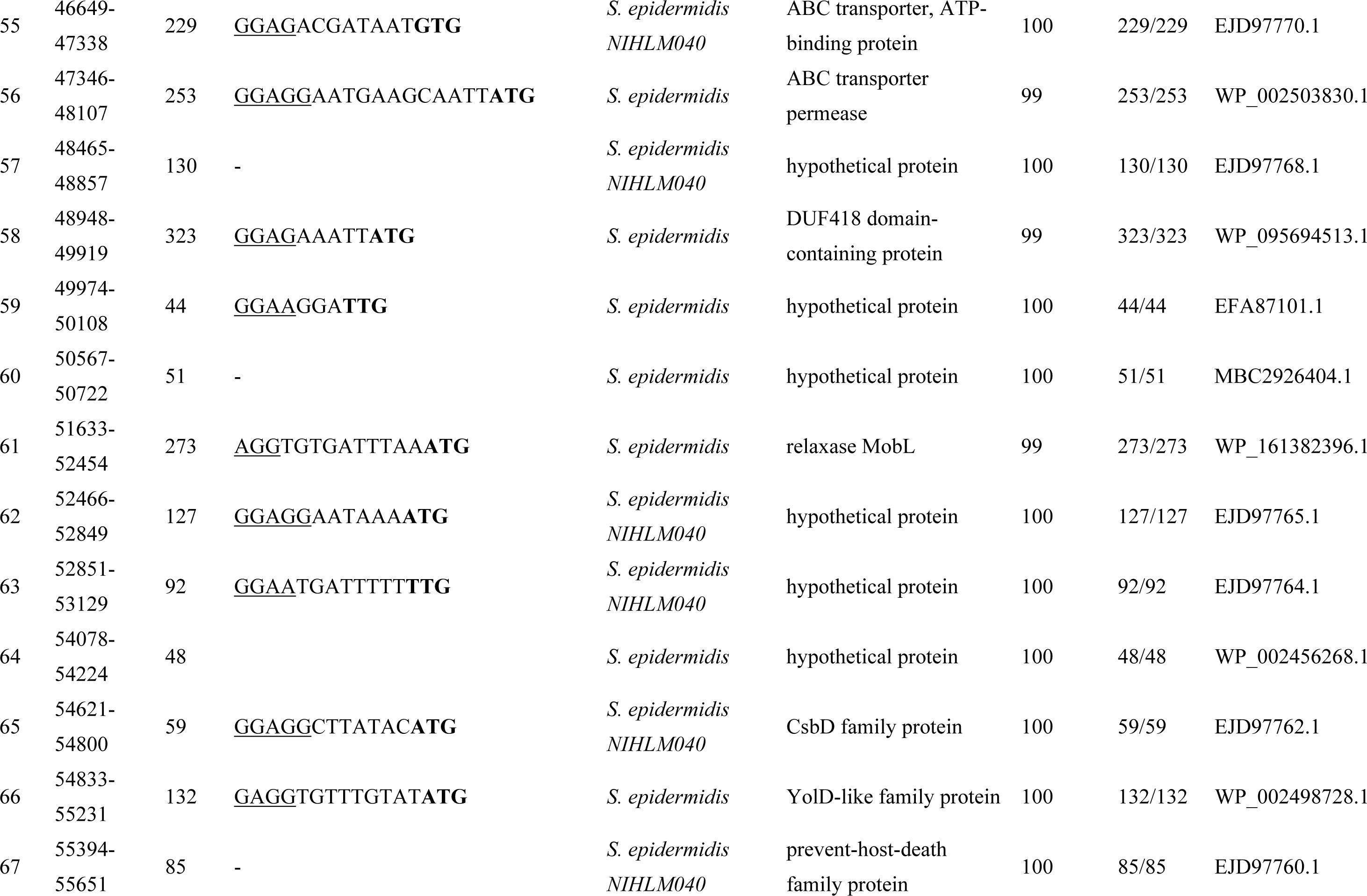
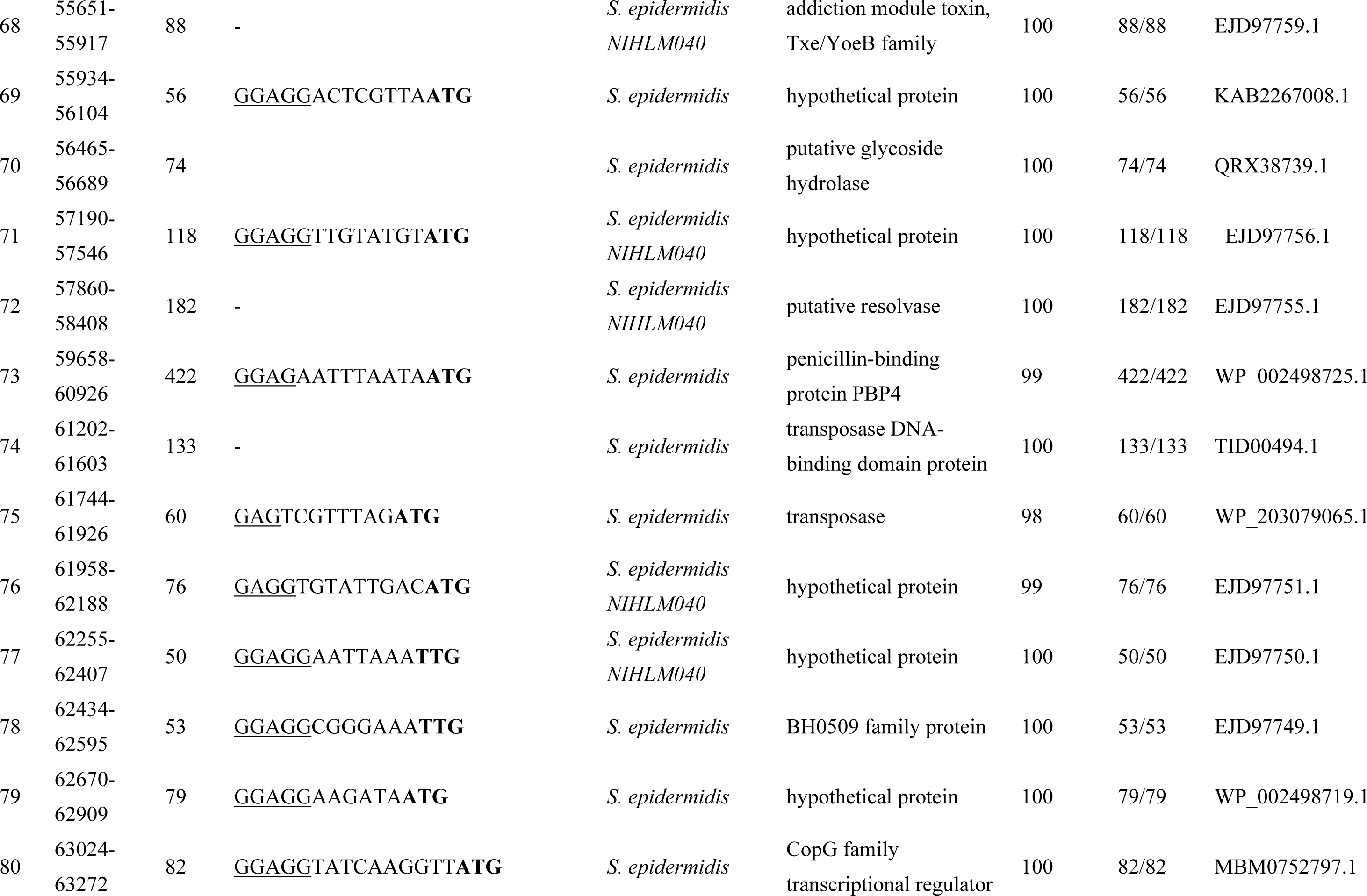
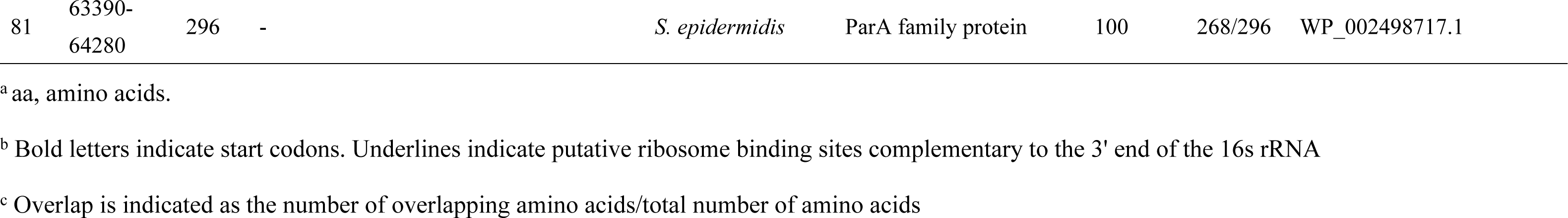
Genes in pEpi56

### Nucleotide sequence of nukacin-encoding plasmid

The size of the entire plasmid, pNuk650, was 26,160 bp, with 29 open reading frames (ORFs). The plasmid contained nukacin KSE650 synthesis genes (*nukA* coding for prepeptide nukacin KSE650, posttranslational modification enzyme genes *nukM*, processing and secretion transporter genes *nukT*, and immunity protein genes *nukFEGH*), replication-related genes, and other genes including genes coding for hypothetical proteins (Fig. 3a, Table 3). Compared to the plasmid pIVK45 (21,840 bp), which carried the gene coding for nukacin IVK45 (16), pNuk650 was larger with a higher number of ORFs (Fig. 3a). The amino acid sequence of nukacin KSE650 showed similarity to nukacin IVK45 with one mismatch at the 4^th^ position, but displayed lower similarity to nukacin ISK-1 with 10 mismatches [36][29] (Fig. 3b). The mature peptide of nukacin KSE650 showed a perfect match with nukacin IVK45 and 5 mismatches with nukacin ISK-1.

**Table. 3.**
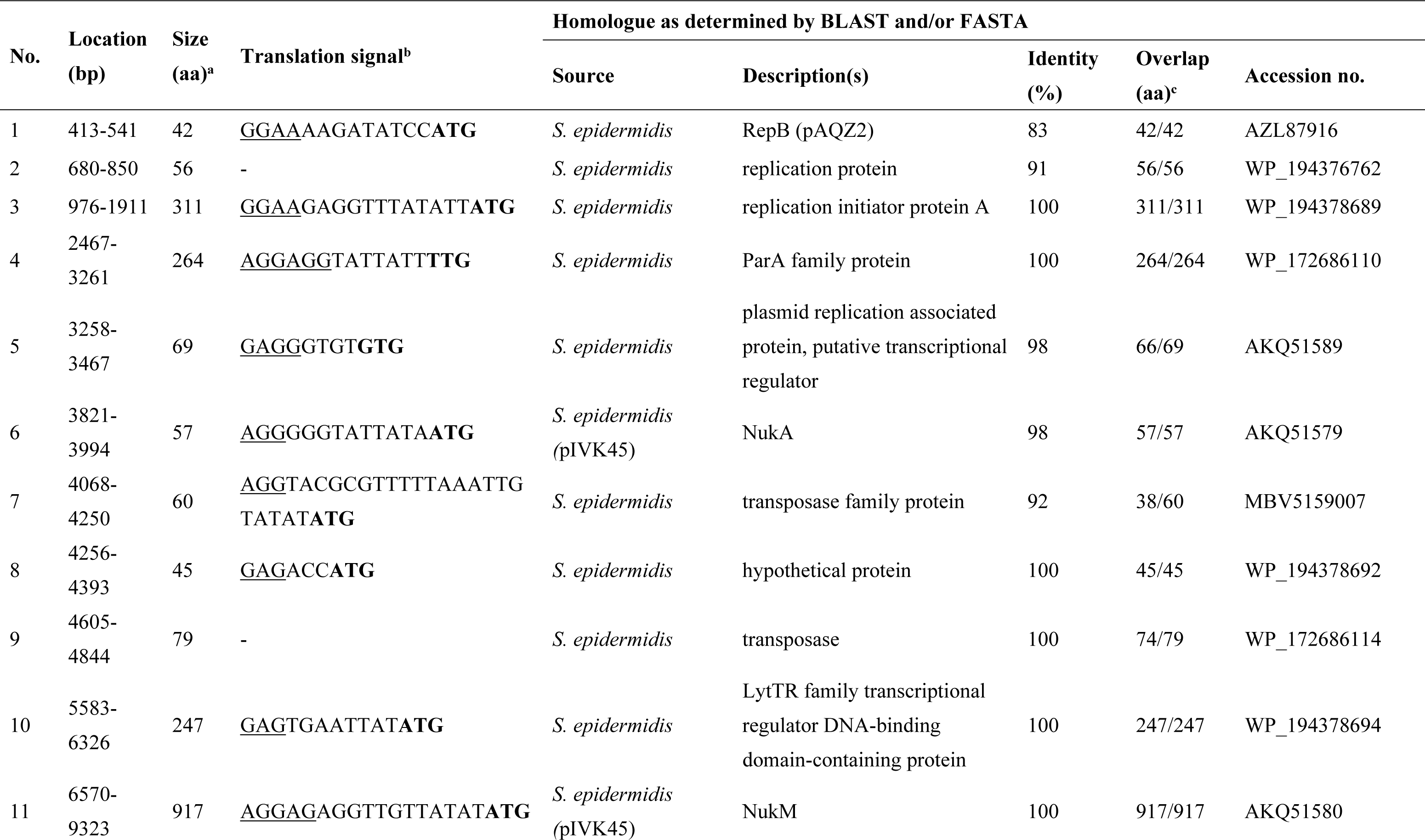
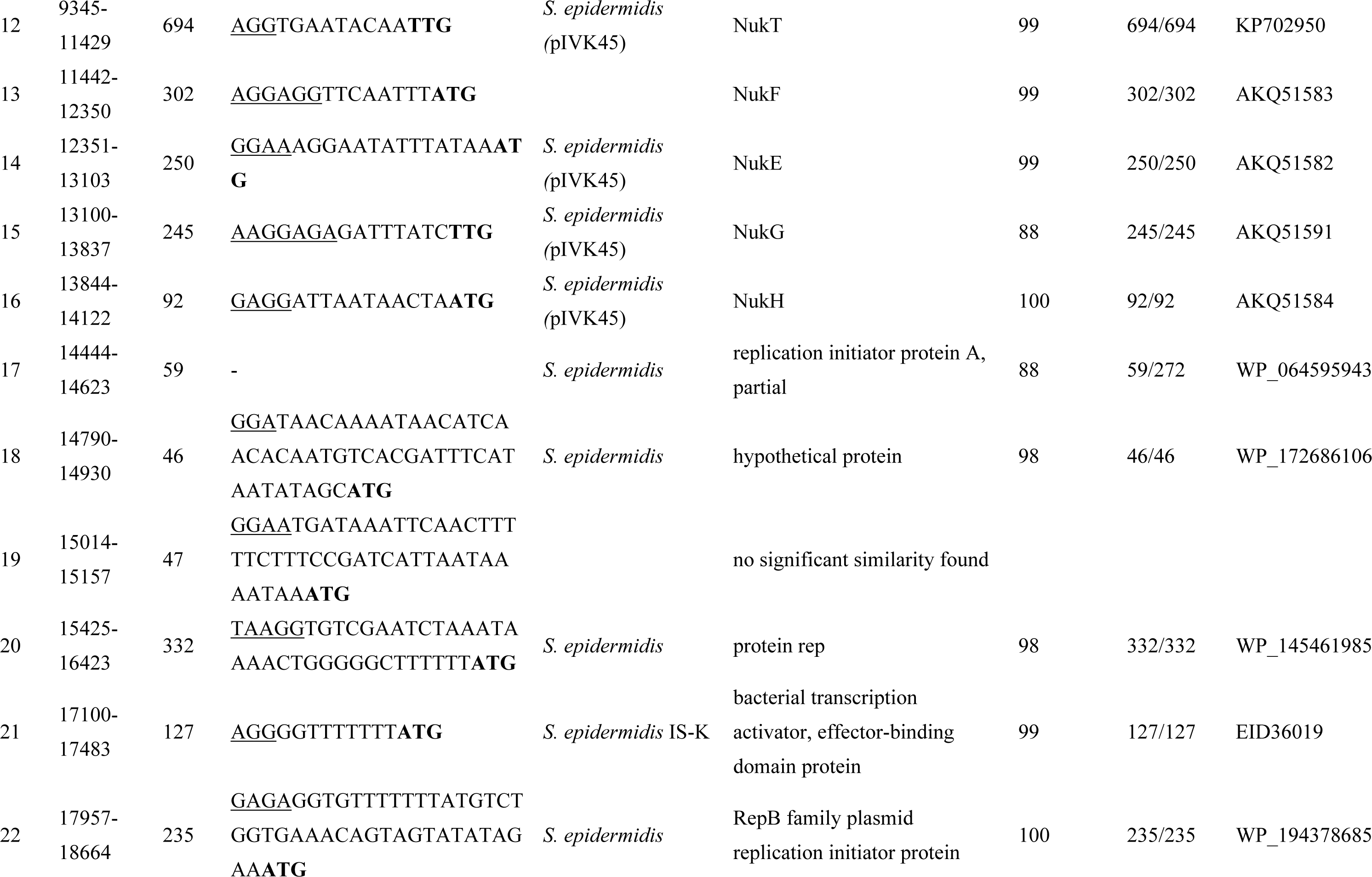
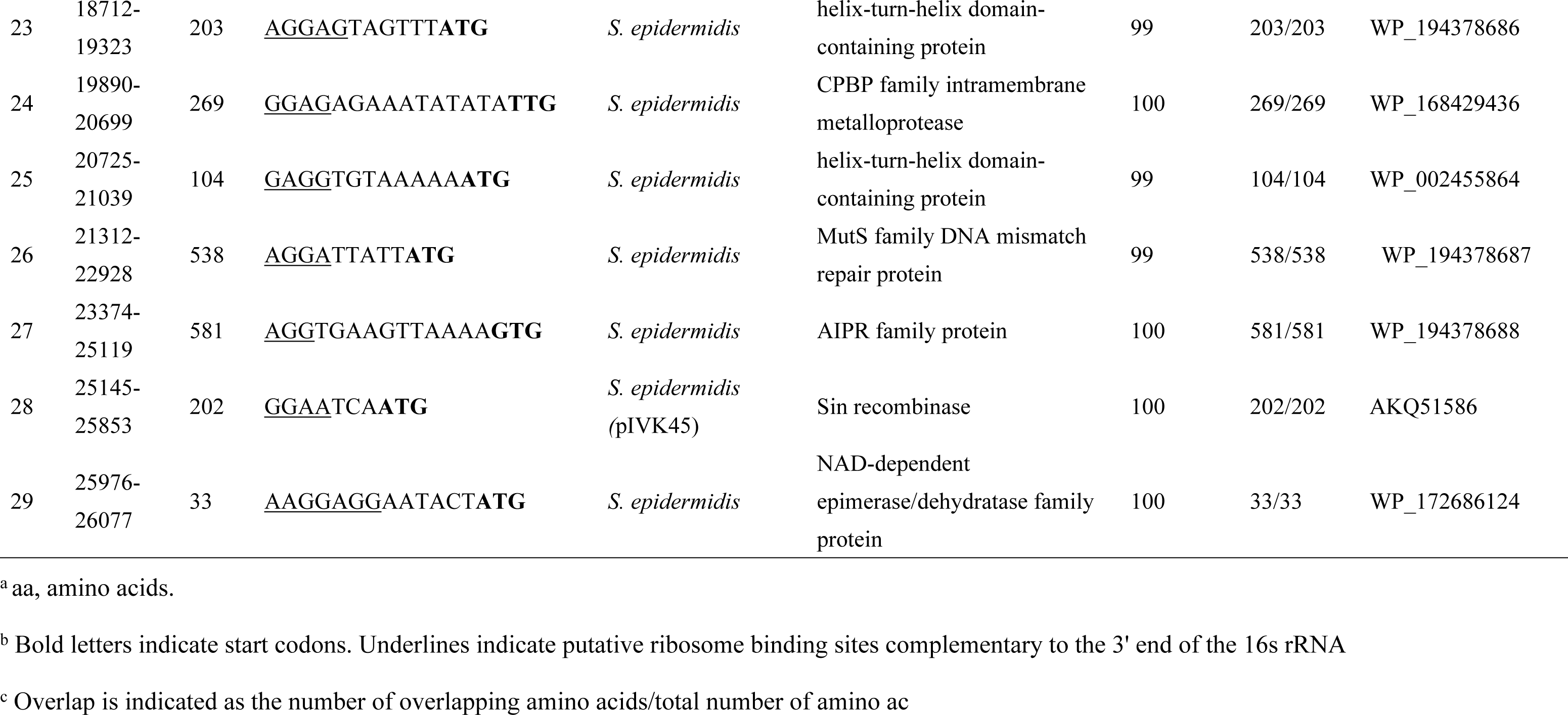
Genes in pNuk650

**Figure 3.**
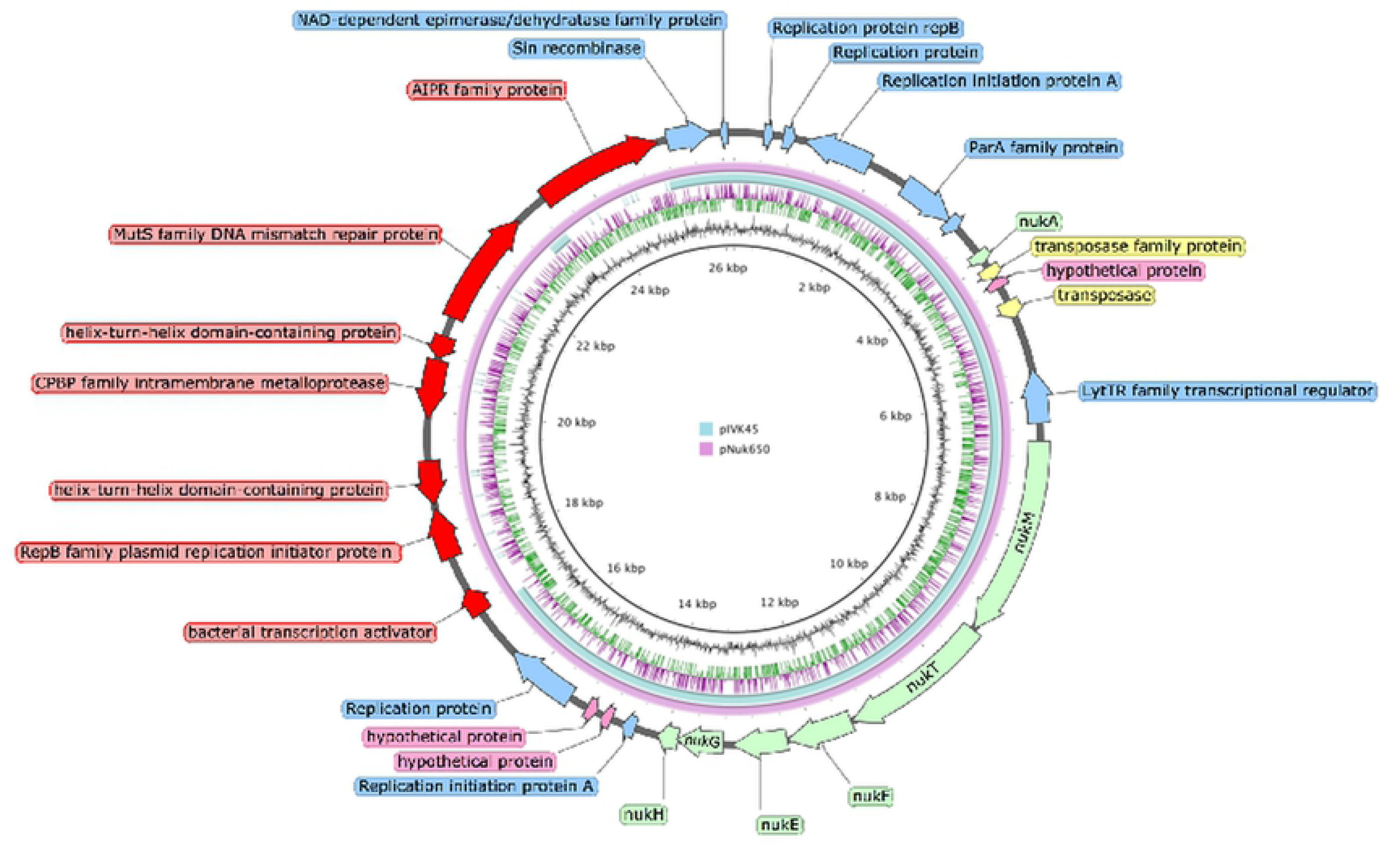
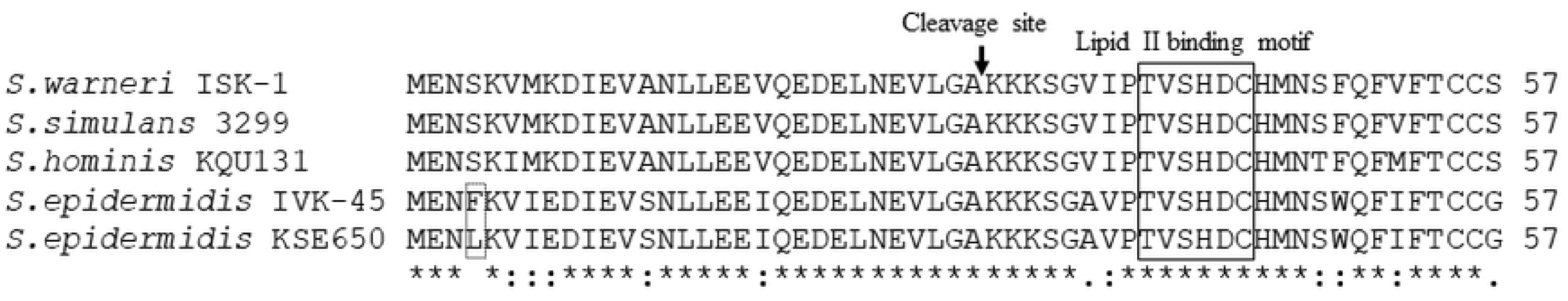
Comparison of the plasmids between *S. epidermidis* KSE650 and IVK45 strains. (a) Nukacin-encoding plasmid from KSE650 (pNuk650) and the comparison with pIVK45 (b) Amino acid alignment of nukacin ISK-1, nukacin 3299, nukacin KQU131, nukacin IVK45 and nukacin KSE650.

### Identification of epidermin KSE56 and nukacin KSE650

Epidermin KSE56 and nukacin KSE650 were purified from the culture supernatant of KSE56 and KSE650, respectively. Using ESI-MS analysis, the molecular masses of purified epidermin KSE56 and nukacin KSE650 were found to be 2163.95 Da and 2938.33 Da, respectively. The mass of these peptides corresponded to calculated mass of epidermin (2163.95 Da) and nukacin KSE650 (2938.33 Da).

### Antibacterial activity of epidermin KSE56 and nukacin KSE650 against several skin and oral commensal bacteria

In this study, *S. epidermidis* strains were isolated from the oral cavity. *S. epidermidis* is also known as a commensal bacterium. Therefore, we investigated the antibacterial activity of the two bacteriocins against oral and skin commensal bacterial species.

We first performed a direct assay using KSE56, KSE650 and plasmid-deleted strains. The plasmid-deleted strains showed no inhibitory zone against *S. hominis*, while the wild-type strains, KSE56 and KSE650, displayed inhibitory zones (Fig. 4).

**Figure 4.**
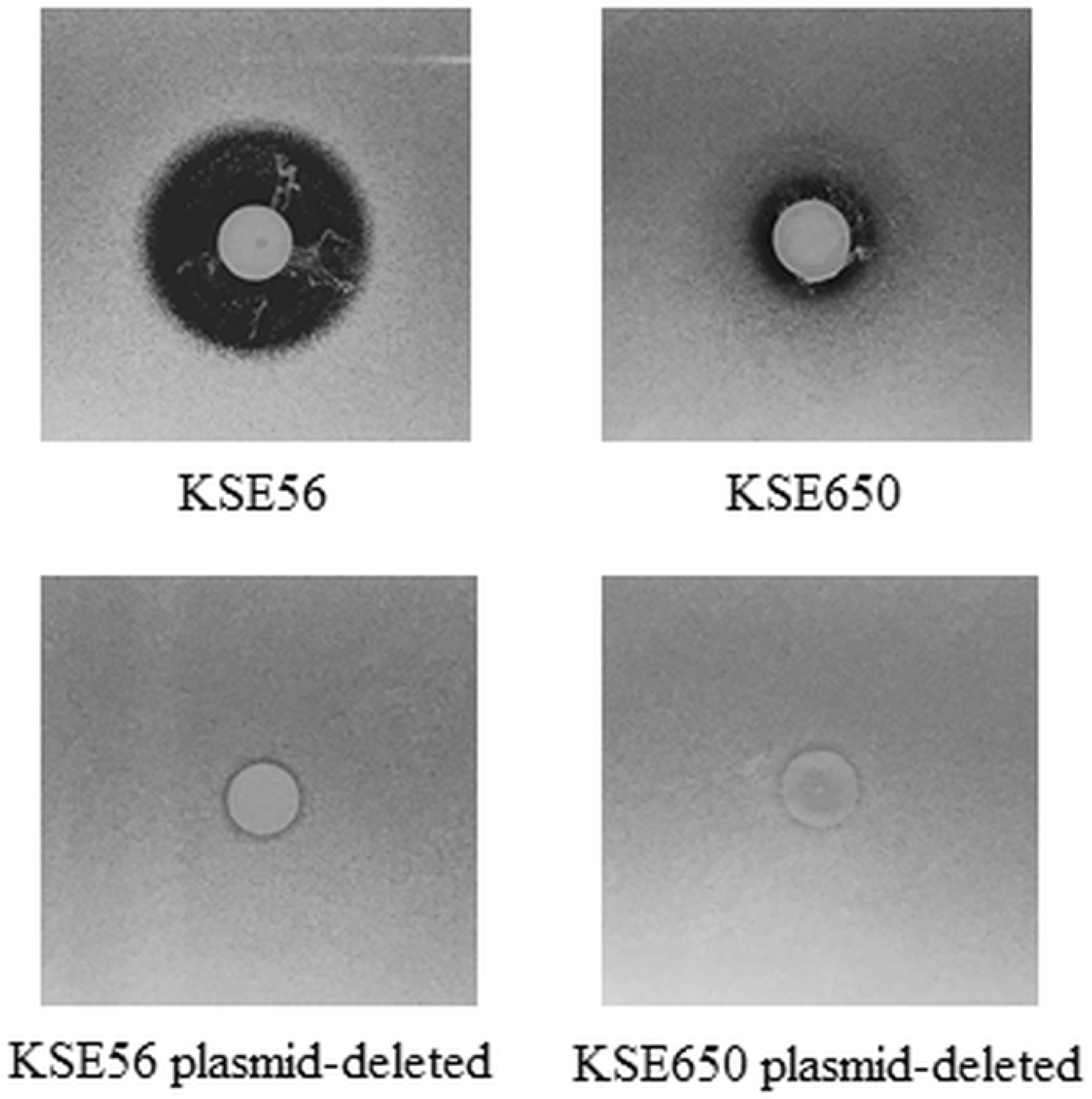
Antibacterial activity of KSE56, KSE650 and their plasmid-deleted strains. Direct assays were performed using KSE56, KSE650 and their plasmid-deleted strains. *S. hominis* was used as an indicator strain.

Afterwards, we performed a direct assay using KSE56 and KSE650 (Table 4). The epidermin-producing strain, KSE56, showed a strong antibacterial activity (>20 mm diameter of inhibitory zone) against *M. luteus*, *C. pseudodiphtheriticum, S. captis*, and *S. hominis*, and an activity (>5 mm diameter) against *R. mucilaginosa*, *S. haemolyticus*, *S. simulans*, and *S. saprophyticus*. KSE56 also showed an antibacterial activity against *S. epidermidis* without bacteriocin production (KSE1, 10, 12, 16), plasmid-curing KSE56 and plasmid-curing KSE650. The inhibitory zone was not observed in *S. epidermidis* KSE56, *S. epidermidis* KSE650, *C. accolens*, the *S. warneri* ISK-1 and *S. aureus* strains. Regarding oral streptococci, KSE56 showed a strong activity against *S. salivarius* and *S. gordonii*, and modest activity against *S. mutans* and *S. sanguinis*.

**Table 4.** Antibacterial activity of KSE56 and KSE650 against various bacterial species

| Indicator strains | Halo size (mm) |  |  |
| --- | --- | --- | --- |
|  | KSE56 | KSE650 | *S. warneri* |
| *Corynebacterium pseudodiphtheriticum* JCM1320 | 10.0±0.8 | 10.7±0.5 | 11.7±0.5 |
| *Corynebacterium accolens* JCM8331 | 5.0 | 5.0 | 11.3±0.5 |
| *Micrococcus luteus* JCM1464 | 31.7±1.2 | 27.0±0 | 33.0±0 |
| *Rothia mucilaginosa* JCM10910 | 8.7±0.5 | 8.0±0 | 13.0±0 |
| *Staphylococcus haemolyticus* JCM2416 | 14.3±0.9 | 13.3±0.5 | 16.0±0.8 |
| *Staphylococcus capitis* JCM2420 | 27.7±0.9 | 27.3±0.5 | 17.3±0.5 |
| *Staphylococcus simulans* JCM2424 | 12.7±0.5 | 28.7±0.5 | 22.7±0.5 |
| *Staphylococcus saprophyticus* JCM20595 | 12.0±1.6 | 12.3±0.5 | 13.3±0.5 |
| *Staphylococcus hominis* JCM31912 | 26.3±0.5 | 16.3±0.5 | 21.7±0.5 |
| *Staphylococcus epidermidis* KSE1 | 12.3±0.5 | 7.0±0.8 | 5.0 |
| *Staphylococcus epidermidis* KSE10 | 12.0±0 | 7.3±0.5 | N.D.<sup>1</sup> |
| *Staphylococcus epidermidis* KSE12 | 17.0±0.8 | 9.7±0.5 | N.D. |
| *Staphylococcus epidermidis* KSE16 | 14.3±0.5 | 8.7±0.5 | N.D. |
| *Staphylococcus epidermidis* KSE56 | 5.0 | 5.0 | 5.0 |
| *Staphylococcus epidermidis* KSE650 | 5.0 | 5.0 | 5.0 |
| *Staphylococcus epidermidis* KSE56 plasmid-deleted | 20.3±0.5 | 11.3±0.5 | N.D. |
| *Staphylococcus epidermidis* KSE650 plasmid-deleted | 11.0±0 | 11.7±0.5 | N.D. |
| *Staphylococcus warneri* ISK-1 | 5.0 | 5.0 | 5.0 |
| *Staphylococcus aureus* MW2 | 5.0 | 5.0 | 11.3±0.5 |
| *Staphylococcus aureus* COL | 5.0 | 5.0 | 11.0±0 |
| *Staphylococcus aureus* RN4220 (MSSA) | 5.0 | 5.0 | 10.7±0.5 |
| *Streptococcus mutans* UA159 | 15.0±0.8 | 5.0 | 5.0 |
| *Streptococcus sanguinis* GTC217 | 12.0±0 | 5.0 | 10.3±0.9 |
| *Streptococcus salivarius* GTC215 | 27.7±0.5 | 12.3±0.5 | 18.3±0.5 |
| *Streptococcus gordonii* JCM12995 | 29.0±0 | 17.0±0 | 23.0±0 |

<sup>1</sup>Not determined

The nukacin KSE650-producing strain KSE650, showed strong antibacterial activity (>20 mm diameter) against *M. luteus*, *S. captis*, and *S. simulans* and activity (>5 mm diameter) against *C. pseudodiphtheriticum ,R. mucilaginosa*, *S. haemolyticus*, *S. hominis*, and *S. saprophyticus*. KSE650 also showed an antibacterial activity against *S. epidermidis* without bacteriocin production (KSE1, 10, 12, 16), plasmid-curing KSE56 and plasmid-curing KSE650. The inhibitory zone was not observed in *S. epidermidis* KSE56, *S. epidermidis* KSE650, *C. accolens*, *S. warneri* ISK-1 and *S. aureus* strains. Regarding oral streptococci, KSE650 showed activity against *S. salivarius*, and *S. gordonii*, and no activity against *S. mutans* and *S. sanguinis*. Compared to the nukacin ISK-1-producing *S. warneri* strain, *S. warneri* showed stronger activity against commensal and oral bacteria except *S. capitis* and *S. simulans*. Notably, *S. warneri* ISK-1 showed activity against the *S. aureus* strain.

Next, we investigated the antibacterial activity of KSE56 and KSE650 against each TCS-inactivated mutant in *S. aureus* (Fig.5). The *apsRS*- and *braRS*-inactivated mutants showed an inhibitory zone compared to the WT and the other TCS- inactivated mutants. In particular, the *braRS*-inactivated mutant showed the strong susceptibility to these strains.

**Figure 5.**
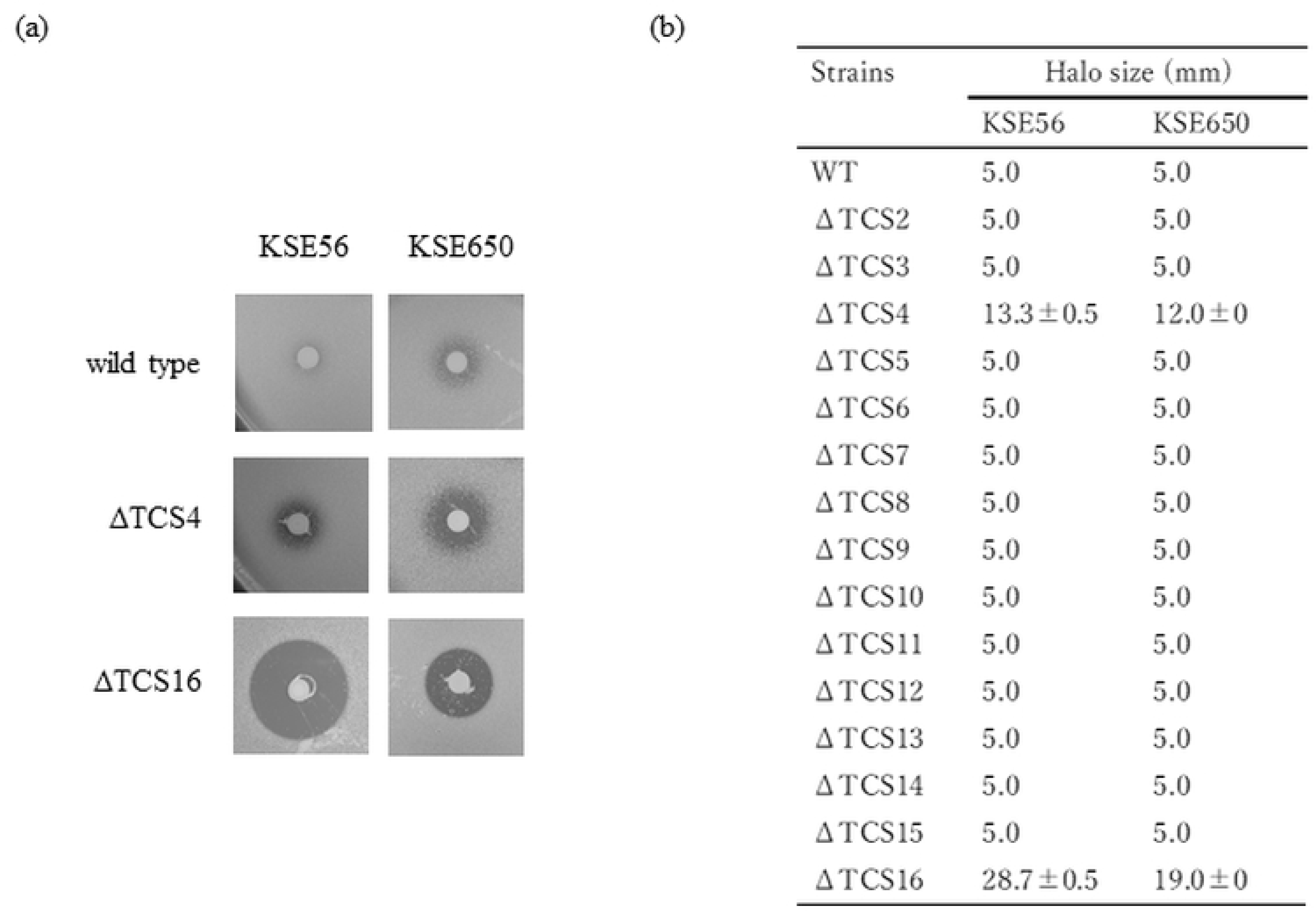
Antibacterial activity of KSE56, and KSE650 against *S. aureus* TCS- inactivated mutants. Direct assay was performed using KSE56 and KSE650. Fourteen sets of TCS- inactivated *S. aureus* mutants were used as indicator strains (a). Three independent experiments were performed. The diameter of the inhibitory zone was measured and the average values were calculated (b).

Co-culture of S. aureus with S. warneri or L. lactis

Cocultures of *S. epidermidis* KSE1 (bacteriocin negative), KSE56, and KSE650 with *M. luteus* were analysed. In coculture with *M. luteus*, the proportion of *S. epidermids* KSE1 was 46.2%, while the proportions of KSE56 and KSE650 were 70.4% and 79.8%, respectively (Fig. 6).

**Figure 6.**
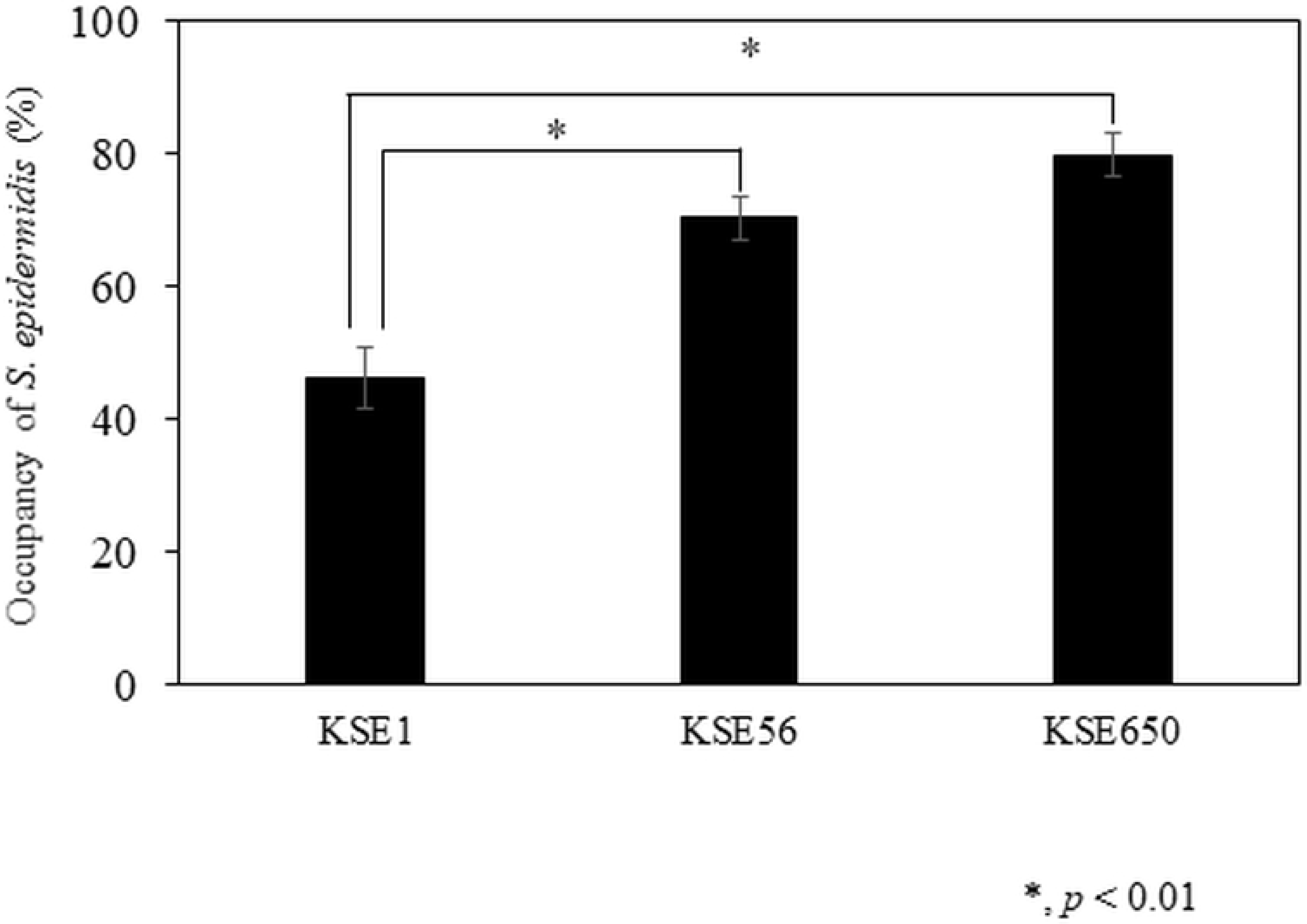
The proportion of *S. epidermidis* KSE1, KSE56 and KSE650 cocultured with *M. luteus*. Coculture assays were performed according to the method described in the Materials and methods. Post hoc multiple comparisons were made using Tukey’s test.

## Discussion

In this study, we tried to isolate *S. epidermidis* strains that produced bacteriocin. We used the *S. aureus* MW2 *braRS*-inactivated mutant as the indicator strain for screening. We previously reported that BraRS was involved in resistance to several bacteriocins including nisin A, nukacin ISK-1 and bacitracin [34]; therefore, a *braRS*-inactivated mutant increased susceptibility to these bacteriocins. Nisin A and nukacin ISK-1 are lantibiotics that act against lipid II molecules, which are responsible for cell wall biosynthesis, and subsequently, form a pour complex[37]. In addition, it was reported that many gram-positive bacteria, including staphylococci, streptococci, bacilli, lactococci and enterococci, produced lantibiotics that bind to lipid II [12,16,38,39,17–24] Therefore, the *braRS*-inactivated mutant is a good indicator strain to screen lipid II-binding lantibiotics. Finally, we identified 2 strains that produce epidermin and nukacin IVK45-like bacteriocins. Whole genome analysis of the 2 strains revealed that both genes were located on the plasmids (Fig. 2a and 3a).

Epidermin was first identified in the *S. epidermidis* Tü3298 strain [16, 40]. In the Tü3298 strain, epidermin is located on the plasmid, pTu32. Recently, the whole genome sequence of the Tü3298 strain was determined [41], but the entire plasmid sequence of pEpi56 was not reported. Therefore, our study is the first to report the complete nucleotide sequence of epidermin harbouring plasmids. Additionally, the epidermin-producing strain identified in this study was the second strain, following the Tü3298 strain. The nucleotide sequence of the *epiA* coding epidermin showed 2 mismatches between the two strains, but the amino acid sequence was similar. When the epidermin synthesis genes were compared between the 2 strains, *epiT* showed a significant difference (Fig. 2b). *epiT* in KSE56 was intact, while this gene in Tu3298 was disrupted into 2 genes, *epiT’* and *epiT’’* in Tü3298.

EpiT is involved in the secretion of the peptide. In previous reports that demonstrated the antibacterial activity of epidermin in Tü3298 [16–18], epidermin was correctly modified and secreted externally. However, Peschel A et al reported that the introduction of intact *gdmT*, encoding the secretion protein for gallidermin, which was close to epidermin in Tü3298, increased the production of epidermin in culture supernatant [42]. Therefore, the secretion activity of epiT’/T’’ is considered to be partial, while the intact *epiT* gene in KSE56 may be responsible for full secretion of the epidermin peptide.

Nukacin IVK-1 was first identified in *S. warneri* [29]. Since then, nukacin ISK-1 like bacteriocins have been identified in *S. epidermidis* [25], *S. hominis* [43], and *S. simulans* [44]. The amino acid sequence of KSE650 shows a high similarity with that of IVK45 by only one mismatch in the entire peptide, and 100% match with the mature peptide. Comparison of the plasmid between the two strains showed that KSE650 was larger than Tü3298, but the composition and the order of nukacin- related genes were identical (Fig. 2a). The larger size of pNuk650 was due to the insertion of an approximately 8 kbp fragment, which was detected in pNuk650 but not in pIVK45 (Fig. 3a, red arrows).

The antibacterial activity of these peptides against skin and oral commensal bacteria (oral streptococci) showed different patterns. In particular, the epidermin- producing strain (KSE56) had antibacterial activity against oral streptococci, while nukacin-producing strains had less activity. Interestingly, comparing nukacin ISK-1 and nukacin KSE650 suggested that 5 amino acid differences (Fig. 7) were responsible for the different activities against several bacteria used in this study.

**Figure 7.**
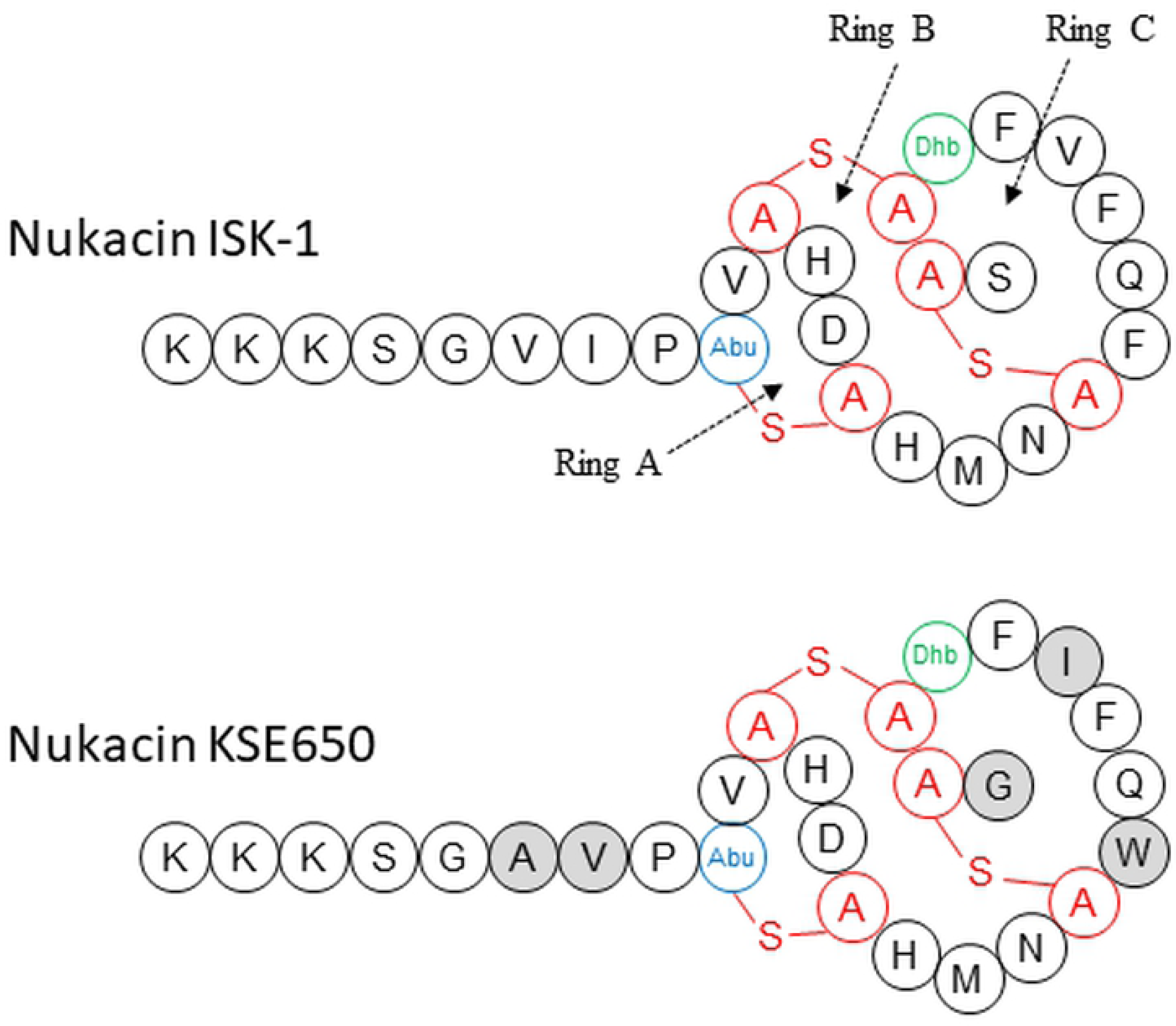
Structure of nukacin ISK-1 and nukacin KSE650. The mature peptide sequences of nukacin ISK-1 and nukacin KSE650 are shown. The deduced calculated mass of mature nukacin KSE650 is consistent with that observed by ESI-MS. The structure is identical to that of nukacin ISK-1, except for the residues indicated by grey circles. Dhb, Ala-S-Ala, and Abu-S-Ala indicate dehydrobutyrine, lanthionine, and 3-methyllanthionine respectively.

Previously, it was reported that the structure of ring A in nukacin ISK-1 binds to the pyrophosphate moiety of lipid II, the precursor for cell wall peptidoglycan biosynthesis, and ring C was also associated with the binding of the isoprene chain [45]. Since lipid II molecules are widely conserved among gram positive bacteria, the different antibacterial activities between nukacin ISK-1 and nukacin KSE650 are influenced by the other molecules specific to each bacterial species. Furthermore, it is noteworthy that epidermin and nukacin KSE650 showed no inhibitory zone against *S. epidermidis* KSE650 and KSE56, respectively, while epidermin and nukacin KSE650 showed an activity against plasmid-curing KSE650 and plasmid-curing KSE56, respectively (Table 4). Although the immunity factors for epidermin and nukacin KSE650 were EpiFEG and NukFEG/NukH, respectively, which could be found in a respective plasmid, our results indicate that these immunity factors showed a cross-resistance to another bacteriocin. We previously reported that BraRS and ApsRS, TCSs, are involved in resistance to nisin A and nukacin ISK-1 [34]. Since *S. epidermidis* also possesses TCSs with similarity to BraRS and ApsRS, *S. epidermidis* TCSs may be involved in the resistance to epidermin and nukacin KSE650.

In conclusion, we determined the complete sequence of two plasmids encoding epidermin and nukacin KSE650 in *S. epidermidis* isolated from the oral cavity. *S. epidermidis* is the major commensal bacterium in human skin and the oral cavity. Based on our findings of the direct assay and coculture assay, it is speculated that bacteriocins produced by *S. epidermidis* affect the bacterial composition of the host flora, including the skin, nasal and oral flora. However, in this study, we focused on the isolation of lantibiotic-producing strains using a *braRS*-inactivated strain as the indicator. Therefore, it is possible that *S. epidermidis* also produces other types of bacteriocins. Further studies are required to demonstrate the influence of *S. epidermidis* bacteriocins on the formation of bacterial flora.

## Acknowledgments

We thank Dr. Tomoko Amimoto, the Natural Science Center for Basic Research and Development (N-BARD), Hiroshima University for the measurement of ESI-MS analysis.

## Author contributions

Conceptualization: Miki Kawada-Matsuo, Norifumi Nakamura, Hitoshi Komatsuzawa

Data curation: Kenta Nakazono, Mi Nguyen-Tra Le

Formal analysis: Kenta Nakazono, Mi Nguyen-Tra Le

Funding acquisition: Miki Kawada-Matsuo, Norifumi Nakamura, Hitoshi Komatsuzawa, Motoyuki Sugai

Investigation: Kenta Nakazono, Mi Nguyen-Tra Le, Noy Kimheang, Junzo Hisatsune, Yuichi Oogai, Miki Kawada-Matsuo

Methodology: Motoyuki Sugai, Miki Kawada-Matsuo, Hitoshi Komatsuzawa

Project administration: Miki Kawada-Matsuo, Norifumi Nakamura, Hitoshi Komatsuzawa

Resources: Masanobu Nakata, Miki Kawada-Matsuo, Norifumi Nakamura, Motoyuki Sugai, Hitoshi Komatsuzawa

Software: Mi Nguyen-Tra Le

Supervision: Masanobu Nakata, Miki Kawada-Matsuo, Norifumi Nakamura, Motoyuki Sugai, Hitoshi Komatsuzawa

Validation: Masanobu Nakata, Miki Kawada-Matsuo, Norifumi Nakamura, Hitoshi Komatsuzawa

Visualization: Masanobu Nakata, Miki Kawada-Matsuo, Hitoshi Komatsuzawa

Writing – original draft: Kenta Nakazono, Mi Nguyen-Tra Le, Miki Kawada-Matsuo

Writing – review & editing: Miki Kawada-Matsuo, Mi Nguyen-Tra Le, Masanobu Nakata, Yuuichi Oogai, Norifumi Nakamura, Motoyuki Sugai, Hitoshi Komatsuzawa

## Supporting information

S1 Fig. Comparison of amino acid sequences of EpiT between the KSE56 and Tü3298 strains

S2 Fig. Comparison of nucleotide (A) and amino acid sequences (B) of *epiA* between the KSE56 and Tü3298 strains

## Notes

### Competing Interest Statement

The authors have declared no competing interest.

